# Metabolic reprogramming and synergistic cytotoxicity of genistein and chemotherapy in human breast cancer cells

**DOI:** 10.1101/2024.11.13.623452

**Authors:** Sandra Tobón-Cornejo, Ariana Vargas-Castillo, Mandy Juarez, Joshua Ayork Acevedo-Carabantes, Lilia G. Noriega, Omar Granados-Portillo, Alma Chávez-Blanco, Rocío Morales-Bárcenas, Nimbe Torres, Armando R. Tovar, Alejandro Schcolnik-Cabrera

## Abstract

Breast cancer (BCa) is a heterogeneous disease, initially responsive to hormone therapy but often developing resistance to both hormonal and chemotherapy treatments. Novel therapeutic strategies are needed for drug-resistant BCa. Genistein, a phytoestrogen structurally similar to estrogen, competes with estrogen for receptor binding and exhibits anti-cancer effects. In this study, we investigated the cellular and metabolic impacts of genistein, alone or in combination with chemotherapy, in two human BCa cell lines—one estrogen receptor-positive (ER+) and one estrogen receptor-negative (ER−). We observed a strong synergistic effect on cell viability at low concentrations of genistein and chemotherapy, resulting in reduced clonogenic capacity and impaired cell migration. Genistein alone modulated cellular energy metabolism, notably reducing ATP production in MCF7 (ER+) cells. This metabolic shift was linked to a decreased dependence on fatty acids for energy, coupled with a decrease in the rate-limiting mitochondrial translocase CPT1 required for fatty acid oxidation, alongside with an increase in intracellular fatty acid levels. While the most significant changes occurred in ER+ cells, ER− cells also showed responses to genistein treatment. Collectively, our findings suggest that low dose genistein, in combination with conventional chemotherapy, induces synergistic anti-cancer effects, promoting cellular senescence. This effect may be partly mediated by a reduced reliance on fatty acid metabolism in BCa cells.

## 1. Introduction

Breast cancer (BCa) is the most prevalent cancer among women, particularly affecting younger females, in whom the disease tends to be highly aggressive [1,2]. In the United States, BCa, along with lung cancer, and colorectal cancer (CRC), accounts for 51% of all new cancer diagnoses in women, with BCa representing ∼32% of these cases. For 2024, an estimated 310,720 new BCa diagnoses are projected [1]. Furthermore, approximately 42,250 deaths are expected to occur due to BCa in 2024 [2]. Early-stage treatment significantly improves survival outcomes for BCa patients [3]; however, nearly 90% of patients with metastatic BCa succumb to the disease due to therapy resistance [4], underscoring an urgent need for novel therapeutic options for BCa.

Recent years have seen an ascending trend in the incidence of BCa, and this trajectory is projected to continue for the coming decades [3]. One of the primary factors driving this rise is the widespread adoption of “Westernized” lifestyles [4]. Notably, up to 40% of all global cancer cases are linked to dietary factors, particularly the chronic consumption of refined sugars, alcohol and animal-derived calories [5, 6]. In contrast, diets rich in phytochemical compounds, such as genistein, have been identified as potential protective factors against cancer development [7, 8].

Genistein is an anti-lipogenic isoflavone that controls the master metabolic regulator AMPK [9], and under obesity and insulin resistance conditions, it has demonstrated to modulate fatty acid oxidation (FAO) as well [10, 11]. In cancer, however, the role of genistein remains controversial: while low concentrations may promote oncogenesis, higher doses exhibit anti-cancer properties [12]. This dual function is attributed to the chemical structure of genistein, which competes with estrogens for binding to the estrogen receptor alpha (ERα), a key target in certain cancer subtypes, including BCa [13]. Additionally, while the chronic consumption of genistein, such as through soy products, is considered safe [14], conventional cancer therapies like chemotherapy are associated with numerous secondary effects [15], which significantly impair quality of life [16] and contribute to mortality in up to 7.5% of treated patients [17]. Consequently, there is a pressing need to reduce chemotherapy doses to mitigate these adverse effects and improve the quality of life for cancer patients.

Genistein plays a key role in modulating lipid metabolism within cells. Oral administration of genistein, in combination with a high-fat diet, has been shown to reduced subcutaneous adipose tissue, lower levels of free fatty acids (FFAs), and decrease low-density lipoprotein, collectively contributing to a reduction in body weight in mouse models [18]. In human colorrectal cancer cells, *in vitro* studies have demonstrated that genistein exerts anti-lipogenic effects by reducing the mRNA expression of 3-hydroxy-methylglutaryl-coenzyme A (HMG-CoA) reductase, a crucial enzyme for the *de novo* synthesis of cholesterol [19]. Conversely, a study using mammary tissue from healthy donors, who later developed BCa, revealed that metabolic rewiring—marked by the upregulation of lipid metabolism and adipogenesis-related genes—was a critical step in the progression toward malignancy [20].

Although the anti-cancer potential of genistein has been previously explored, its interactions with clinically used chemotherapy, and the associated metabolic outcomes in BCa, remains unclear. In this study, we aimed to investigate in two human BCa cell lines, one estrogen receptor-positive (ER+) and one estrogen receptor-negative (ER-). Specifically, we evaluated how these drug combinations influenced anti-cancer activity and metabolic process in a synergistic way, with a particular focus on the anti-lipogenic properties of genistein.

## 2. Materials and methods

### 2.1 Cell culture

The human BCa cell lines MCF7 and MDA-MB-238 (ATCC), positive and negative for the ER, respectively, were employed in this study. Cells were plated in DMEM/F12 (Gibco) medium supplemented with 10% fetal bovine serum (Corning) and 1% streptomycin/amphotericin (Gibco) (complete medium), at 37 °C, in a 5% CO_2_ incubator.

### 2.2 Drugs

Genistein (Enzo), tamoxifen (Invitrogen), and docetaxel (Taxoterete), were dissolved in DMSO (Sigma) for all the assays. The compounds were administered alone, or in the GT (genistein+tamoxifen) or GD (genistein+docetaxel) schemes.

### 2.3 Cell viability and identification of inhibitory concentrations (ICs)

5x10^4^ cells/well were seeded in 6-well plates (Costar), with 2 mL of complete medium. After 24h of pre-incubation, cells were treated during 72h with increasing concentration of each compound alone or their controls, composed of the vehicle at the same volume as of that of the highest evaluated concentration. Fresh complete medium containing each drug/vehicle was changed every 24h. After 72h, cells were counted as previously described [21]. The cytotoxic effect was expressed as the percentage of cell viability relative to control cells. The percentage of growth inhibition was estimated with the SigmaPlot software V10.0 (Systat Software, Inc.), and the IC_10_-IC_50_ values were determined.

### 2.4 Pharmacological interaction

Increasing concentrations of genistein (IC_10_, IC_20_, IC_30_, IC_40_, IC_50_) were combined with their respective increasing concentrations of either tamoxifen (IC_10_, IC_20_, IC_30_, IC_40_, IC_50_) or docetaxel (IC_10_, IC_20_, IC_30_, IC_40_, IC_50_), for MCF7 and MDA-MB-231, respectively. The resulting mixed solutions were incubated for 72h in 5x10^4^ cells/well, as indicated above. The pharmacological interaction was determined with the combination index (CI) method from the formula of Chou and Talalay, using the Calcusyn software V2.0 (Biosoft) [22]. The synergistic combination at the lowest concentration was selected for further experiments.

### 2.5 Clonogenic assays

Cells were treated during 72h as indicated above. Next, 1x10^3^ living cells/condition were recovered and plated in new 6-well plates for clonogenic assays, as described by Juarez M. *et al*. [21]. The resulting colonies were counted 5 days after seeding using ImageJ V2.0 (NIH, MA, USA).

### 2.6 Wound-healing assay

4.5x10^4^ MCF7 or 6x10^4^ MDA-MB-231 cells/well were seeded in 6-well plates, as indicated above. After 24h of pre-incubation, 90% of confluence was ensured and cells were washed once with 1X PBS. Next, a 200-μL pipette tip was used to do a straight scratch in each well, and cells were treated for 24h with the conditions referred above. Images of the wounds at times 0h and 24h were photographed with a digital camera (Leica DMC2900), and the percentage of wound closure was calculated using ImageJ.

### 2.7 Oxygen consumption and extracellular acidification rates

Oxidative phosphorylation, glycolysis and FAO were assessed by quantifying the oxygen consumption (OCR) and extracellular acidification rates (ECAR) with a Seahorse Bioscience Extracellular Flux Analyzer XF96e (Seahorse Bioscience). Briefly, 4x10^3^ cells/well were seeded into XF96 culture microplates (Seahorse Bioscience) with 100μL complete medium, and 24h after incubation they were treated for 72h. For each condition, 5 wells were tested. 1h prior to each Seahorse assay, cells were equilibrated with bicarbonate-free low buffered medium (Seahorse Bioscience), pH 7.4, without supplements, at 37°C in a non-CO_2_ incubator. All required reagents for each experiment were prepared in Seahorse assay medium and loaded into the cartridges for 30 min into Seahorse plates as the experiment progressed. All results were normalized by cell number using a BioTek Cytation 1 equipment where the cells were previously stained with 2µM Hoechst (Invitrogen). Mitochondrial respiration and glycolysis were determined with the Mito Stress and Glycolysis Stress Test Kits, as previously reported [23, 24]. Palmitate oxidation indicative of lipid oxidation was performed with the Palmitate Oxidation Stress Test Kit, according to Nasci et al [25]. Normalized results were analyzed using the Wave software (Seahorse Biosciences).

### 2.8 Fuel flexibility

OCR was measured to assess effects on mitochondrial respiration using a Seahorse XFe96 Analyzer (Agilent). Following the 72-hour treatment period, metabolic capacity and dependency analysis were performed in treated cells using the XF Mito Fuel Flex Test Kit (Agilent). All results were normalized to the number of cells, and cells were quantified with 2µM Hoechst (Invitrogen) in a BioTek Cytation 1 as referred above. Normalized results were analyzed using the Wave software (Seahorse Biosciences).

### 2.9 Western blot

For total protein extraction, cells and tissues were homogenized using lysis buffer (0.5% sodium deoxycholate, 0.1% SDS, 0.006% sodium azide) plus cOmplete Mini (Roche) and PhosSTOP (Roche). Twenty micrograms of extracted protein were loaded onto SDS-PAGE gels (10%) and transferred to a polyvinylidene difluoride membrane (PVDF) (Bio-Rad). Blocking was undergo using 5% fat-free milk powder, and membranes were then incubated with the following primary antibodies: FAS H300 (1:1000; Santa Cruz, cat. no. sc-20140), α-AMPK H-300 (1:1500; Santa Cruz, cat. no. 25792), α-pAMPK T-172 (1:500; Santa Cruz, cat. no. 33524), α-CPT1 8F6AE9 (1:2000; Abcam, cat. no. 128568), α-SREBP1 C-20 (1:500; Santa Cruz, cat. no. 366), α-GADPH G-9 (1:300 Santa Cruz, cat. no. 365062). The secondary antibodies were, as follows: anti-goat (1:10,000 Abcam, cat. no. 6885), anti-mouse (1:10,000; Abcam, cat. no. 6789), and anti-rabbit (1:10,000; Abcam, cat. no. 6721). Detection was performed according to Tobón-Cornejo S. *et al*. [26].

### 2.10 Fatty acids analysis

FAs were extracted as described by Folch *et al*. [27]. After being extracted, FAs were methylated as previously described [28]. Methylated FAs were analyzed by gas chromatography (Agilent 6850; Agilent, Santa Clara, CA, USA) with a flame ionization detector (Agilent) using a DB-225MS capillary column (J&W Scientific, Albany, CA, USA).

### 2.11 Senescence-associated β-galactosidase assay

To analyze cellular senescence, a senescence-associated β-galactosidase staining kit (Cell Signaling Technology, cat. no. 9860) was used, according to the manufacturer’s instructions. 5x10^4^ cells/well were seeded and treated for 72h, as indicated above. Next, they were washing several times with ice-cold 1X PBS and were fixed for 15 min with 1mL of fixative buffer (Cell Signaling Technology, cat. no. 11674) at room temperature. Finally, cells were dyed with 1mL staining solution (Cell Signaling Technology, cat. no. 11675) and incubated overnight at 37 °C, following the manufacturer’s instructions. Images were captured using a light microscope, and both cell quantification and raw intensity measurement of the dye were conducted in a BioTek Cytation 1.

### 2.12 Statistical analyses

Unless otherwise specified, all experiments were independently performed in triplicate, with three internal replicates. Statistical analyses were performed as follows: drug curves, one-way analysis of variance (ANOVA) with Dunnett post-hoc test; clonogenicity and scratch assays, one-way ANOVA with Tukey post-hoc; Seahorse experiments, one-way ANOVA with Fisher’s LSD post-hoc test; western blot densitometry, chromatography measurements, and senescence-associated β-galactosidase assay, one-way ANOVA with Tukey post-hoc. The results were analyzed with GraphPad Prism V9 (GraphPad, San Diego, CA, USA). Data were expressed as means ± SEM. *p*<0.05 was considered statistically significant.

## 3. Results

### 3.1 Genistein enhances chemotherapy efficacy by synergistically reducing cell viability in breast cancer

Our first objective was to determine the IC_10-50_ values for genistein in the human BCa cell lines MCF7 and MDA-MB-231, positive and negative for ER, respectively. Genistein reduced cell viability in a concentration-dependent manner in both cell lines (**Fig. 1A, 1B, 1D** and **1E**), but the effect was more pronounced in MDA-MB-231 cells (**Fig. 1F**), where lower IC_10-50_ values were observed compared to MCF7 cells (**Fig. 1C**). After identifying the IC_10-50_ values for chemotherapy agents commonly used in BCa treatment tamoxifen for MCF7 and docetaxel for MDA-MB-231 (**Table 1**), we assessed the pharmacology interaction between these agents and genistein. As expected, the combination of genistein+tamoxifen (GT) in MCF7, and of genistein+docetaxel (GD) in MDA-MB-231, resulted in a more significant reduction in cell viability than either drug alone.

**Fig. 1.**
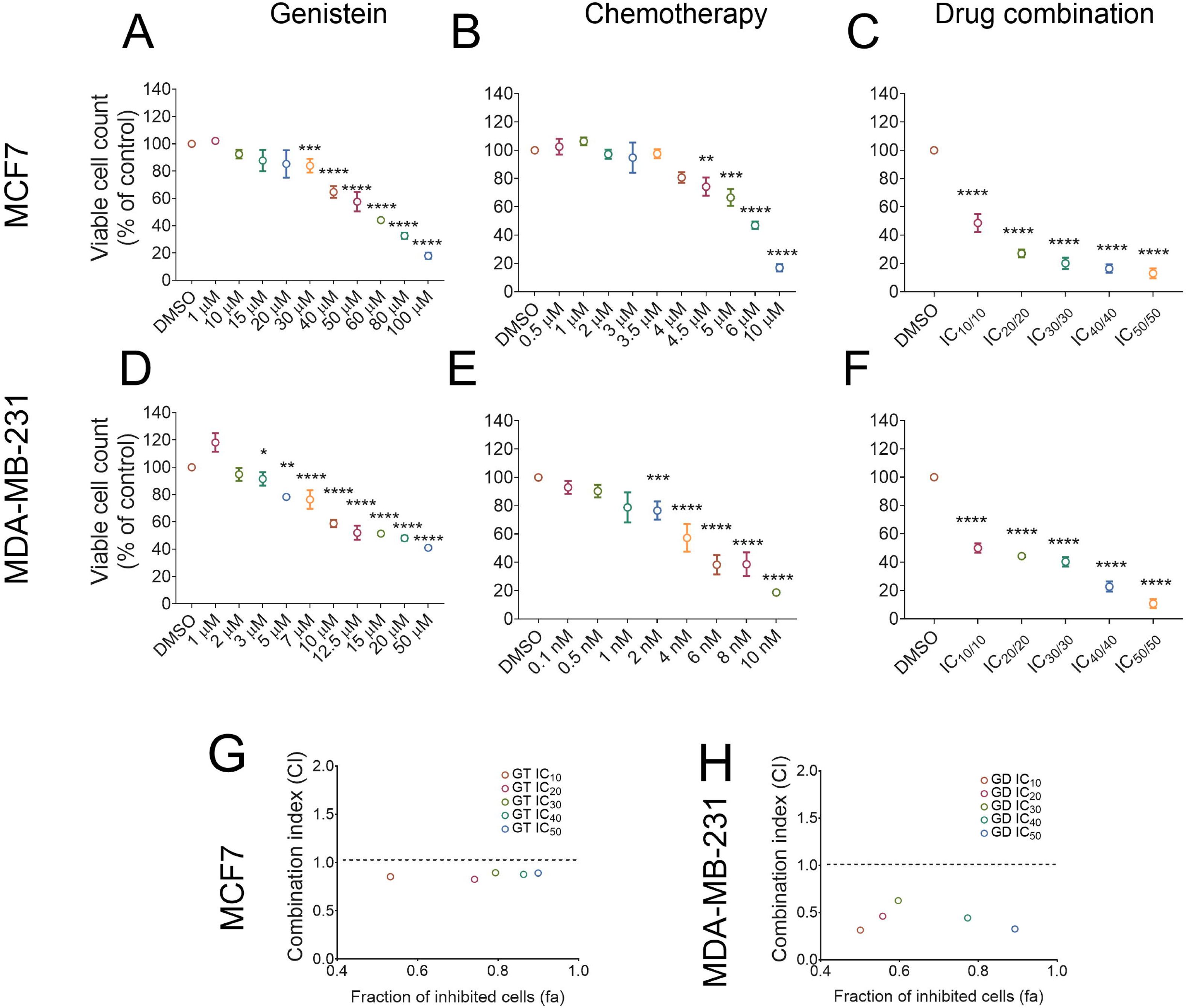
The combination of genistein and chemotherapy synergistically reduces cell viability in breast cancer. Cell viability in MCF7 cells treated with (A) genistein, (B) tamoxifen, and the combination of genistein and tamoxifen (C). Cell viability in MDA-MB-231 cells treated with (D) genistein, (E) docetaxel, and the combination of genistein and docetaxel (F). Combination indexes of the ICs of genistein and tamoxifen in MCF7 cells (G), and of genistein and docetaxel in MDA-MB-231 cells (H)*. All the viability values were evaluated 72 hours after treatment with the referred conditions. Data are presented as mean ± SEM (n=3). Statistical analyses were performed using one-way ANOVA followed by Dunnett’s post-hoc test. *p < 0.05; **p < 0.01; ***p < 0.001; ****p < 0.0001*.

**TABLE 1.** IC concentration per compound and cell line. *GT: Genistein+tamoxifen; GD: Genistein+docetaxel; T: Tamoxifen; D: Docetaxel; IC: Inhibitory concentration*.

| *Cell line* | *Drug scheme* | *Genistein*<br>*concentration* | *Chemotherapy*<br>*concentration* |
| --- | --- | --- | --- |
| ***MCF7*** | **GT IC<sub>10</sub>** | 15.69 µM | 3.62 µM (T) |
|  | **GT IC<sub>20</sub>** | 26.47 µM | 4.26 µM (T) |
|  | **GT IC<sub>30</sub>** | 36.45 µM | 4.78 µM (T) |
|  | **GT IC<sub>40</sub>** | 46.69 µM | 5.29 µM (T) |
|  | **GT IC<sub>50</sub>** | 57.47 µM | 5.83 µM (T) |
| ***MDA-MB-231*** | **GD IC<sub>10</sub>** | 3.53 µM | 0.38 nM (D) |
|  | **GD IC<sub>20</sub>** | 5.02 µM | 1.22 nM (D) |
|  | **GD IC<sub>30</sub>** | 7.04 µM | 2.30 nM (D) |
|  | **GD IC<sub>40</sub>** | 10 µM | 3.59 nM (D) |
|  | **GD IC<sub>50</sub>** | 16.07 µM | 5.05 nM (D) |

Next, we calculated the combination index (CI) to evaluate the nature of the interaction. The CI compares the effects of each drug alone with those of the combination. Values >1, =1, and <1 indicate antagonism, additive effects, and synergy, respectively. Our results showed a strong synergistic effect with the GT and GD combinations, particularly in MDA-MB-231 cells with the GD combination (**Fig. 1G-1H**). This synergism was evident starting at the IC_10_ concentrations of both genistein and chemotherapy (**Table 2**). Based on these findings, we proceeded to test the combination at the lowest concentrations that demonstrated synergy, selecting the IC_10_ values of each drug for both MCF7 and MDA-MB-231 cell lines in subsequent assays.

**TABLE 2.** Drug interaction per cell line. GT: *Genistein+tamoxifen; GD: Genistein+docetaxel; IC: Inhibitory concentration; CI: Combinatory index*.

| *Cell line* | *Drug scheme* | *CI value* |
| --- | --- | --- |
| ***MCF7*** | **GT IC<sub>10</sub>** | 0.854 |
|  | **GT IC<sub>20</sub>** | 0.827 |
|  | **GT IC<sub>30</sub>** | 0.895 |
|  | **GT IC<sub>40</sub>** | 0.878 |
|  | **GT IC<sub>50</sub>** | 0.892 |
| ***MDA-MB-231*** | **GD IC<sub>10</sub>** | 0.314 |
|  | **GD IC<sub>20</sub>** | 0.461 |
|  | **GD IC<sub>30</sub>** | 0.626 |
|  | **GD IC<sub>40</sub>** | 0.443 |
|  | **GD IC<sub>50</sub>** | 0.326 |

### 3.2 The combination of genistein and chemotherapy inhibits cancer cell clonogenic capacity and migration

After 72h of treatment with genistein, chemotherapy, or their combination, 1x10^3^ cells from the MCF7 and MDA-MB-231 cell lines were harvested and re-seeded in new wells to assess clonogenic potential. In MCF7 cells, no significant differences in colony number were observed between the control, genistein, and tamoxifen treatments (**Fig. 2A** and **2B**). However, the GT combination led to a 50% reduction in colony formation. In contrast, MDA-MB-231 cells showed a more pronounced response, with genistein alone reducing proliferation by over 50%, while docetaxel and the GT combination completely inhibited clonogenic capacity (**Fig. 2C** and **2D**).

**Fig. 2.**
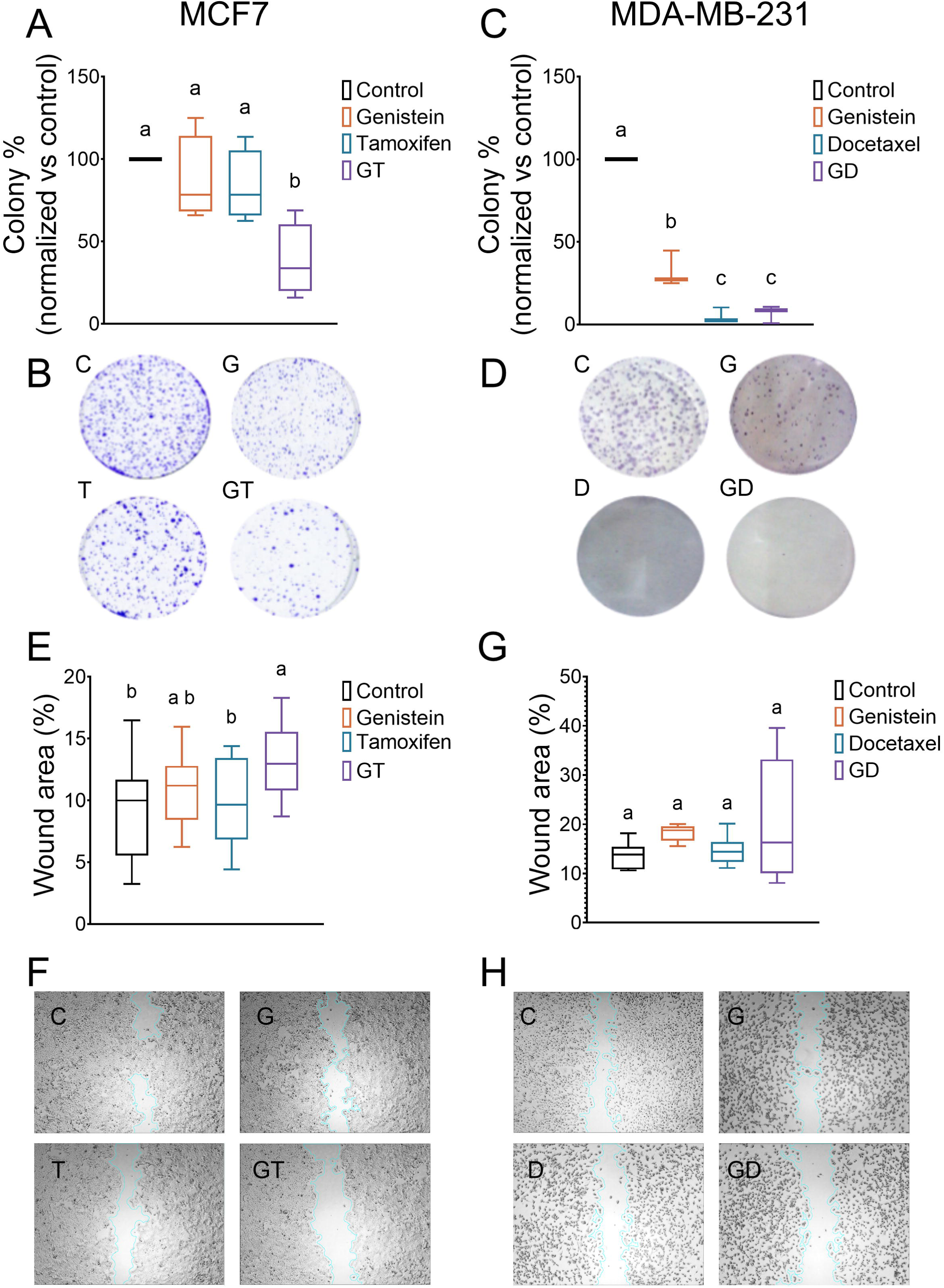
The combination of genistein and chemotherapy reduces the clonogenic capacity and cell migration in breast cancer. Box and whiskers plot showing the percentage of colony formation 5 days after 72 hours of treatment with the referred conditions in MCF7 (A) and in MDA-MB-231 cells (C). Representative plates of colonies stained with crystal violet in MCF7 (B) and MDA-MB-231 cells (D) after 5 days in culture. Box and whiskers plot of the wound area percentage 24h after treatment with the referred conditions in MCF7 (E) and in MDA-MB-231 cells (G). Representative images of the wounds 24h after treatment in MCF7 (F) and in MDA-MB-231 cells (H). *C: control, G: IC_10_ Genistein concentration, T: IC_10_ tamoxifen concentration, GT: combination of IC_10_ genistein and of tamoxifen concentration; D: IC_10_ docetaxel concentration; GD: combination of IC_10_ genistein and of docetaxel concentration. Data are presented as means ± SEM (n=3). Statistical analyses were performed by one-way ANOVA followed by Tukey’s post-hoc test. Multiple comparisons are summarized with lowercase letters (a > b > c > d)*.

Further, we evaluated cellular migration using a wound-healing assay. In MCF7 cells, the GT combination resulted in a significantly lower migration rate, as indicated by a larger unoccupied area in the wound-healing assay (**Fig. 2E** and **2F**). In MDA-MB-231 cells, however, no significant differences were observed in migration between the control, genistein, or docetaxel treatments (**Fig. 2G** and **2H**).

### 3.3 Genistein, alone or in combination with chemotherapy, reduces ATP production and alters the energetic phenotype in MCF7 cells

Given that genistein is a potent metabolic modulator, we next investigated its effects on cellular metabolism using Seahorse assays to assess glycolysis and oxidative phosphorylation. In glycolysis assays, the ECAR revealed no significant reduction in glycolysis, glycolytic capacity, or non-glycolytic acidification in MCF7 cells (**Fig. 3C**). Similarly, tamoxifen treatment did not induce significant changes compared to the control. In contrast, in MDA-MB-231 cells, most glycolytic parameters were downregulated by either genistein or the GD combination, but a significant reduction was only observed in the glycolytic reserve with genistein alone (**Fig. 3D**).

**Fig. 3.**
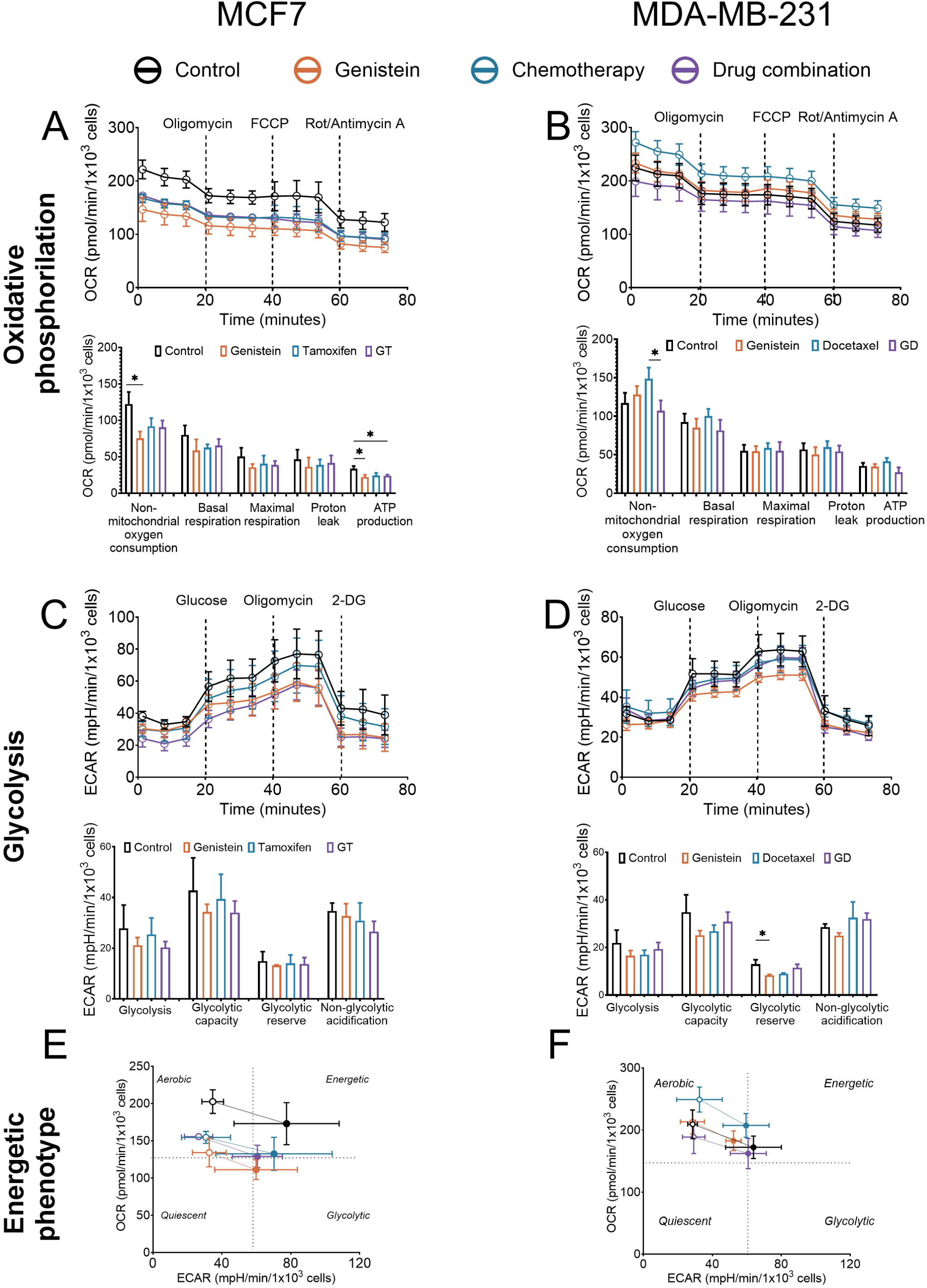
Genistein, alone or in combination with chemotherapy, reduces ATP production and alters the energetic phenotype in MCF7 cells. The oxygen consumption rate (OCR) and individual parameters for mitochondrial respiration in MCF7 cells are shown (A and B); extracellular acidification rate (ECAR) and individual parameters for glycolysis (C and D); and an energetic phenotype chart showing OCR and ECAR under basal conditions and after maximal stress induced by FCCP (E and F) in MCF7 and MDA-MB-231 cells, respectively. *All the values in A-F were evaluated 72h after treatment with the referred conditions. GT: combination of IC_10_ values for genistein and tamoxifen; D: IC_10_ value for docetaxel; GD: combination of IC_10_ values for genistein and docetaxel. Data are presented as means ± SEM (n=4). Statistical analyses were performed using one-way ANOVA followed by Fisher’s LSD post-hoc test. *p < 0.05*.

ATP production was significantly reduced in MCF7 cells treated with either genistein alone or the GT combination (**Fig. 3A**). However, most of the remaining parameters, including basal respiration, maximal respiration, and proton leak, were not significantly altered by these treatments. As with the glycolysis assays, no significant changes were observed when MCF7 cells were treated with tamoxifen alone. In MDA-MB-231 cells, only non-mitochondrial oxygen consumption was reduced with the GD combination compared to docetaxel (**Fig. 3B**). Importantly, when comparing glycolysis and oxidative phosphorylation, more pronounced alterations were observed in oxidative phosphorylation, particularly in MCF7 cells. These findings suggest that genistein alters cellular energy metabolism by impacting glycolysis, FAO, and oxidative phosphorylation, thereby reducing the cell’s capacity to efficiently produce ATP.

### 3.4 Genistein alone does not alter metabolic fuel preference in breast cancer cells

Given the changes in ATP production, particularly in MCF7 cells, we next investigated whether genistein affects the oxidation of specific mitochondrial fuels. Mitochondrial fuel flexibility assays were conducted to measure OCR in the presence of inhibitors targeting different fuel pathways: UK5099 (glucose oxidation inhibitor), BPTES (glutamine oxidation inhibitor), and etomoxir (FAO inhibitor). In MCF7 cells, genistein alone did not significantly alter fuel dependence or capacity for any of the three fuels (**Fig. 4A**). However, the GT combination reduced FA dependence and increased glucose dependence, with no changes observed in glutamine dependence or capacity (**Fig. 4B**). In MDA-MB-231 cells, genistein did not affect the dependence or capacity for any of the fuels tested (**Fig. 4B**). However, docetaxel treatment reduced glucose dependence while increasing both glutamine and FAO capacity (**Fig. 4B**).

**Fig. 4.**
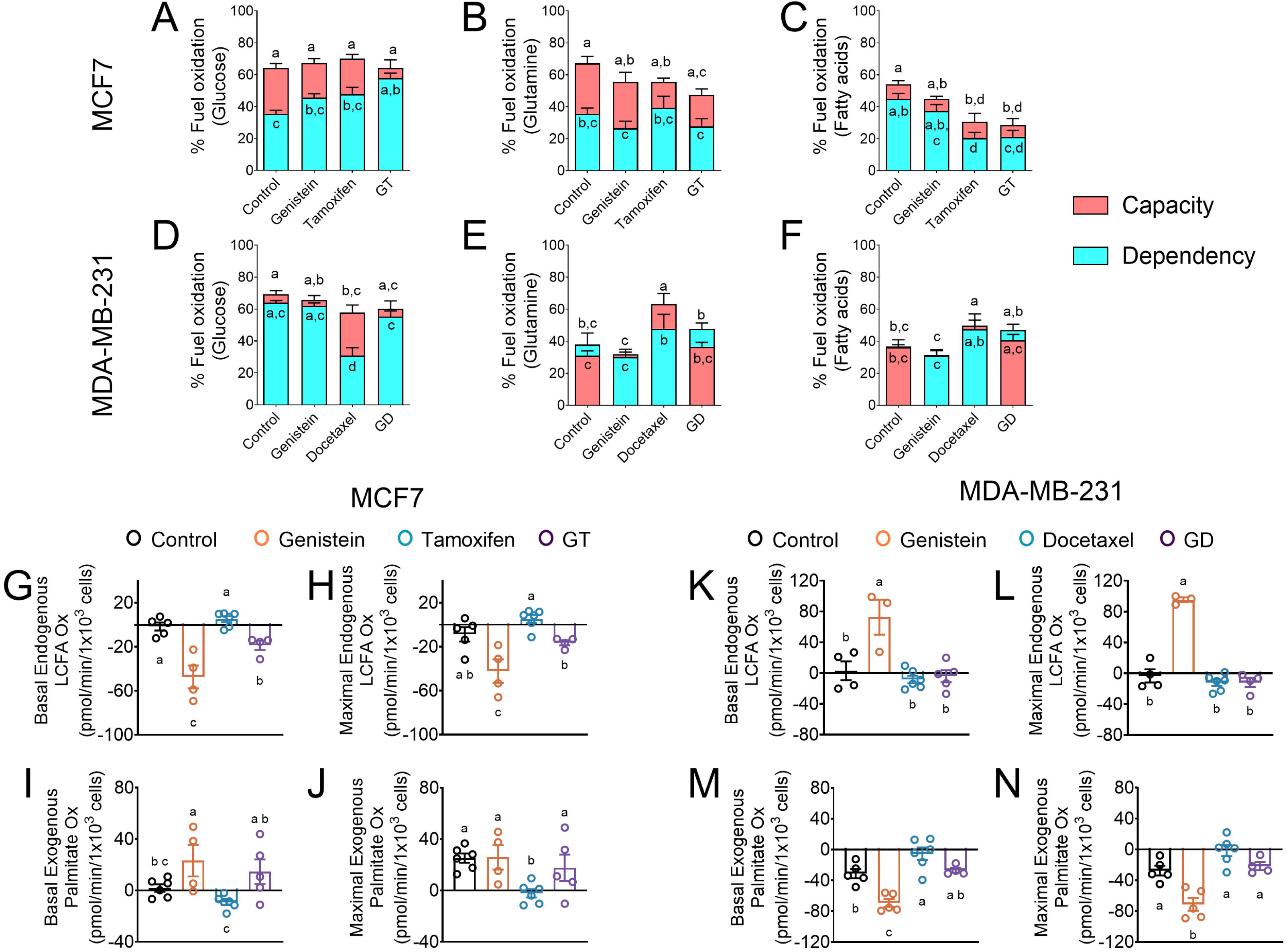
The combination of genistein and chemotherapy reduces the dependency on fatty acids and the endogenous lipid oxidation. Fuel oxidation diagrams represent both the capacity and dependency in MCF7 and MDA-MB-231 cells towards three energetic substrates, named glucose (A and D), glutamine (B and E), and fatty acids (C and F), in basal conditions. (G) Basal and (H) maximal endogenous LCFA oxidation in MCF7 cells. (I) Basal and (J) maximal exogenous palmitate oxidation in MCF7 cells. (K) Basal and (L) maximal endogenous LCFA oxidation in MDA-MB-231 cells. (M) Basal and (N) maximal exogenous palmitate oxidation in MDA-MB-231 cells. LCFA: long chain fatty acids*. All the values in G-N were evaluated 72h after treatment with the referred conditions. GT: combination of IC_10_ values for genistein and tamoxifen; D: IC_10_ docetaxel concentration; GD: combination of IC_10_ values for genistein and docetaxel. Data are presented as means ± SEM (n=3). Statistical analyses were performed by one-way ANOVA followed by Fisher’s LSD post-hoc test. Multiple comparisons are summarized with lowercase letters (a > b > c > d)*.

Collectively, these results suggest that while genistein alone does not significantly alter fuel utilization in either cell line, it can synergistically modify fuel dependency and capacity when combined with chemotherapy in MCF7 cells.

### 3.5 Genistein treatment modifies lipid oxidation in breast cancer cells

Given the significant changes observed in the fuel flexibility assays, we next evaluated lipid oxidation using the Palmitate Oxidation Stress Test to assess both endogenous and exogenous lipid oxidation. In MCF7 cells, treatment with genistein, either alone or in combination with tamoxifen, decreased both basal and maximal endogenous mitochondrial FAO (**Fig. 4G** and **4H**), while increasing basal exogenous lipid oxidation (**Fig. 4G**). In contrast, in MDA-MB-231 cells, genistein alone increased both basal and maximal endogenous mitochondrial FAO (**Fig. 4K** and **4L**). This effect was partially attenuated when genistein was combined with docetaxel (GD), and no significant changes were observed in basal or maximal exogenous lipid oxidation (**Fig. 4M** and **4N**).

These findings suggest that genistein modulates lipid oxidation differently in ER-positive and ER-negative breast cancer cells, with distinct effects on both endogenous and exogenous lipid metabolism.

### 3.6 Genistein modifies the expression of carnitine palmitoyltransferase 1 (CPT1), while simultaneously increasing the intracellular concentration of fatty acids in MCF7 cells

Given the improved alterations in fatty acid (FA) utilization across different treatment schemes in breast cancer cells, we evaluated the expression of key genes involved in lipid metabolism and the intracellular concentrations of FAs. In MCF7 cells, CPT1 expression decreased following genistein treatment while no significant changes were observed with tamoxifen or the combined treatment (**Fig. 5A**). The analysis of intracellular FAs levels revealed that palmitic and stearic acid concentrations increased exclusively with genistein treatment (**Fig. 5C and 5D**). In contrast, oleic acid was elevated both with genistein alone and in combination with tamoxifen (**Fig. 5E**). Arachidonic acid concentrations remained unchanged under all treatment conditions (**Fig. 5F**).

**Fig. 5.**
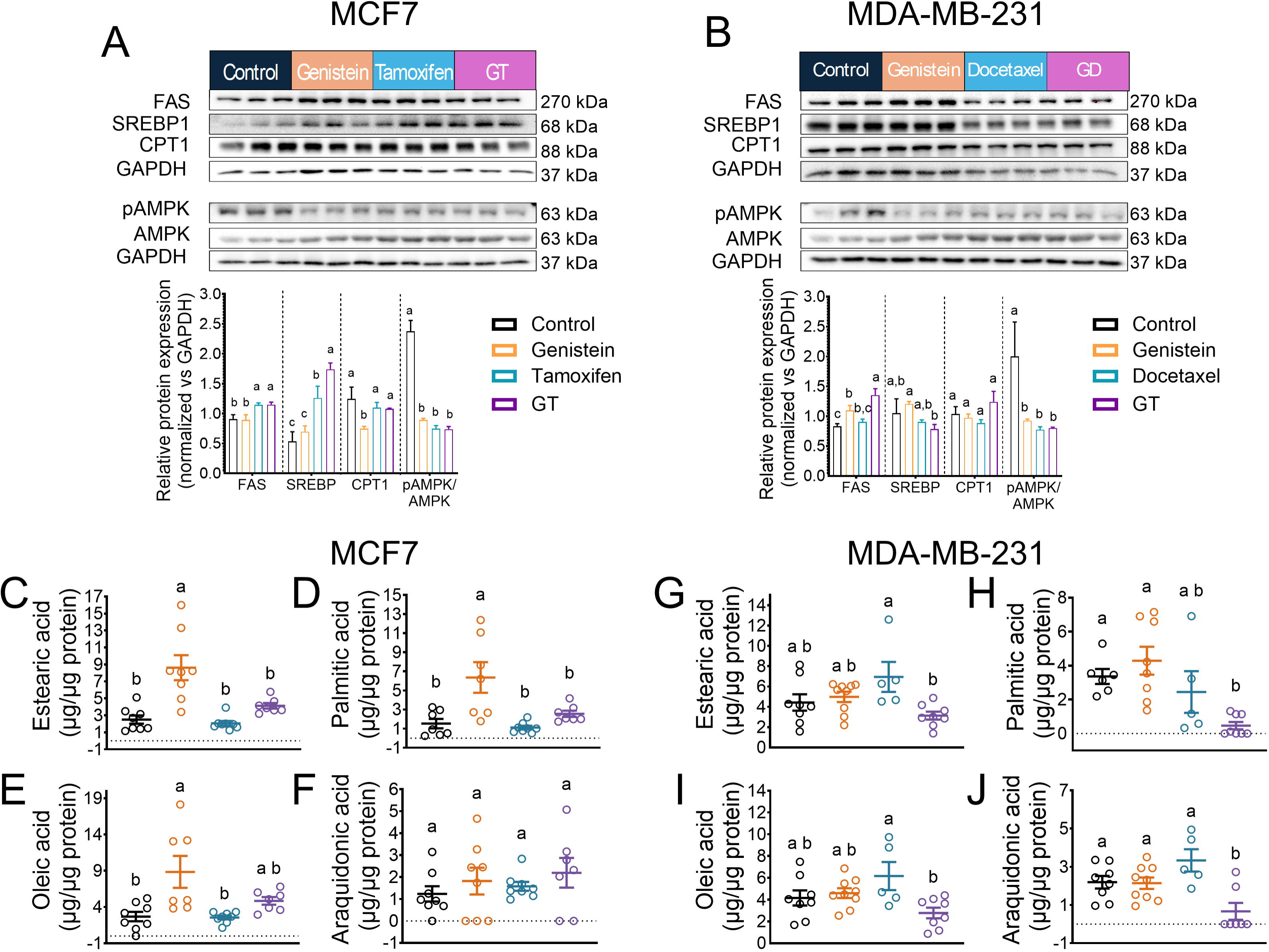
Genistein increases the amount of intracellular fatty acids and reduces the abundance of CPT1 in MCF7 cells. Western blots (top) and their corresponding densitometry analysis (bottom) of FAS, SREBP, CPT1, pAMPK, AMPK, and pAMPK/AMPK, for MCF7 (A) and MDA-MB-231 cells (B) 72h after treatment with the referred conditions. (C-F) Fatty acid concentration in MCF7 cells. (G-J) Fatty acid concentration in MDA-MB-231 cells. *Data are presented as means ± SEM (n=3; each band on the western blots corresponds to a representative independent assay). Statistical analyses were performed by one-way ANOVA followed by Tukey post-hoc test. Multiple comparisons are summarized with lowercase letters (a > b > c > d)*.

In MDA-MB-231 cells, CPT1 expression remained stable (**Fig. 5B**), and no significant changes in palmitic or stearic acid levels were observed (**Fig. 5G** and **5H**). However, a significant decrease in oleic and arachidonic acids concentrations was noted only with the combination of genistein and docetaxel (**Fig. 5I** and **5J**). Regarding AMPK activation, a marked decrease was observed in both breast cancer cell lines under all treatment schemes, suggesting a diminished reliance on FAs as an energy source (**Fig. 5A** and **5B**). This decreased AMPK activity may also be linked to the reduced ATP production observed in previous experiments, particularly in MCF7 cells. Interestingly, in MCF7 cells, treatment with tamoxifen or with its combination with genistein resulted in an increased expression of SRBP1 and FAS (**Fig. 5A**). This may reflect a compensatory response to the reduced FA utilization, potentially aiming to maintain lipid biosynthesis pathway.

### 3.7 The combination of genistein and chemotherapy increases cellular senescence in breast cancer cells

Since the inhibition of proliferation is a hallmark of cellular senescence [29], we investigated whether changes in FA utilization, particularly in MCF7 cells, could affect cell viability by inducing senescence. No significant changes in senescence were observed in cells treated with genistein or tamoxifen alone compared to control (**Fig. 6A** and **6B**). However, the GT scheme led to a 30% increase in senescent cells. A similar trend was observed in MDA-MB-231 cells, where no significant changes were found between control and cells treated with genistein or docetaxel alone (**Fig. 6C** and **6D**). However, the GD combination resulted in a 15% increase in senescent cells. These findings suggest that genistein synergizes with chemotherapy to enhance the inhibition of proliferation through the induction of senescence in both MCF7 and MDA-MB-231 cells.

**Fig. 6.**
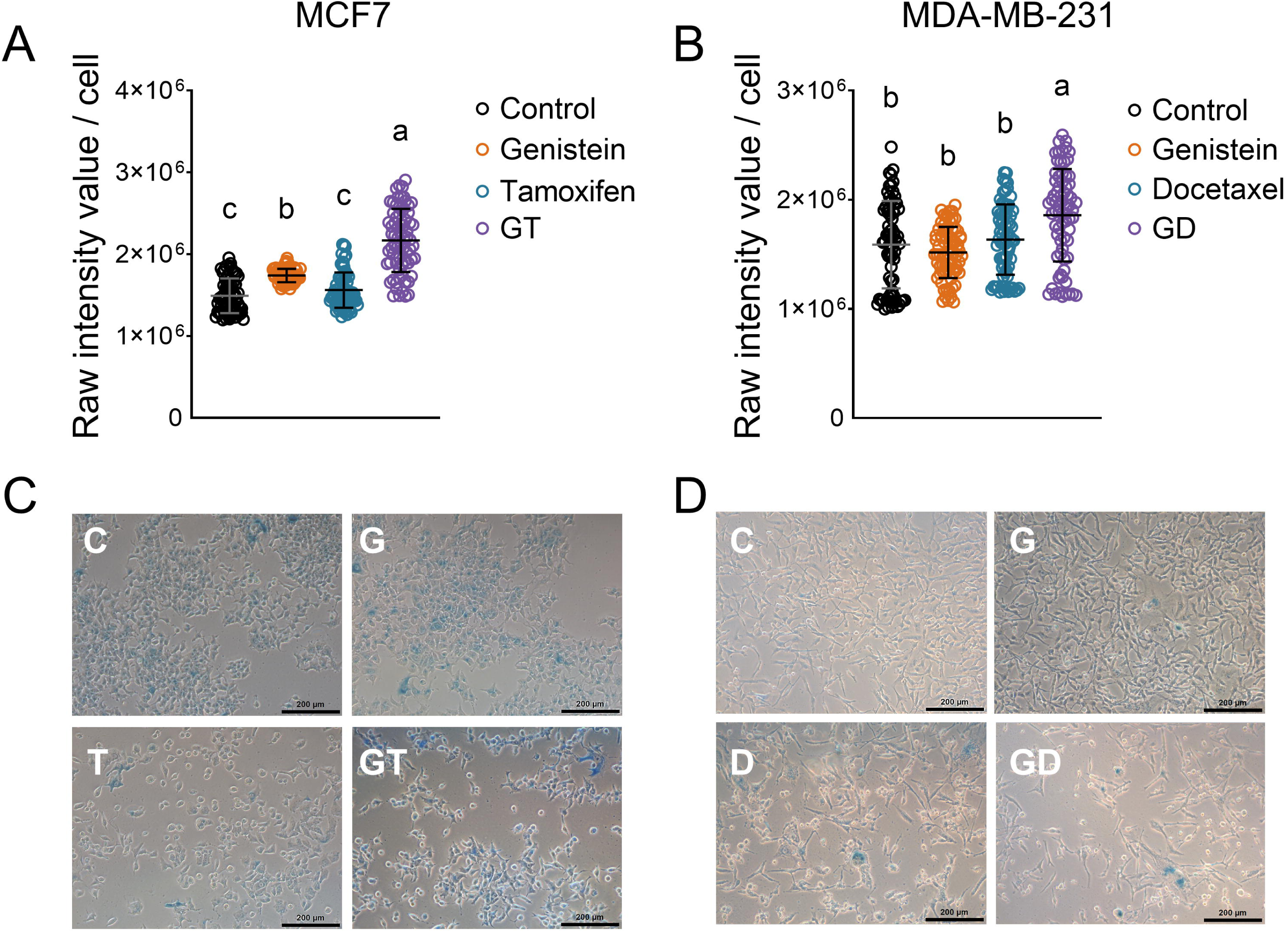
The combination of genistein and chemotherapy induces senescence in breast cancer cells. Intensity values of the senescence-associated SA-β-gal blue staining in MCF7 (A) and MDA-MB-231 cells (B) 72h after treatment with the referred conditions. Representative images of the senescence-associated SA-β-gal blue staining in MCF7 (C) and MDA-MB-231 cells (D). *C: control; G: IC_10_ values for genistein; T: IC_10_ values for tamoxifen; GT: IC_10_ values for genistein and tamoxifen; D: IC_10_ values for docetaxel; GD: IC_10_ values for genistein and docetaxel. Data are presented as means ± SEM. Statistical analyzes were performed by one-ANOVA followed by Tukey post-hoc test. Multiple comparisons are summarized with lowercase letters (a > b > c > d)*.

## 4. Discussion

BCa is the most commonly diagnosed cancer in women worldwide [30]. This high prevalence has driven extensive research aimed at characterizing the disease, identifying potential therapeutic targets, and developing novel treatments tailored to its various subtypes. Given that metabolic abnormalities are now recognized as a hallmark of cancer, identifying specific dysregulated metabolic pathways in malignancies is crucial for uncovering new, targeted therapeutic strategies. This approach is particularly relevant for breast cancer cells, where such metabolic alterations can provide actionable targets for treatment.

In this study, we demonstrate the feasibility of combining the isoflavone genistein with selected chemotherapy agents to synergistically target both ER-positive and ER-negative BCa cells. Isoflavones are natural compounds that function as hormonal modulators and reactive oxygen species scavengers. Among them, genistein is the most biologically active and extensively studied isoflavone [31], known for its anti-cancer properties. Genistein exerts multiple effects, including the activation of caspase-3 and an increase in the BAX/BCL2 ratio—two key events necessary for apoptosis induction. Additionally, genistein inhibits critical signaling pathways, including PI_3_K/AKT, ERK1/2, and MAPK, thereby impairing cellular energy production. It also downregulates the expression of matrix metalloproteinase 9 (MMP-9), limiting the migratory and metastatic potential of cancer cells [32].

In BCa cells, genistein is known to inactivate the HER2 receptor, a proto-oncogene in the epidermal growth factor receptor family, in both ER-positive and -negative BCa cells [33]. The ER-positive MCF7 cell line has been shown to be particularly susceptible to genistein, especially within its stem cell compartment. Fan P. *et al*. reported 32.5μM as the IC_50_ value of genistein for MCF7 [34], which is similar to the IC_50_ value observed in our study in the same cell line. Regarding the ER-negative MDA-MB-231 BCa cell line, 25μM of genistein significantly increases the transcription of *ERα*, rendering the cells more susceptible to receptor-targeted therapies [35]. The latter concentration of genistein can be achieved through dietary supplementary or consumption of soybean-based products [35]. Notably, we have demonstrated that genistein, at similar concentrations, exerts both anti-cancer and anti-lipid effects in BCa cells.

Our results demonstrate that relatively low concentrations of genistein can effectively reduce the cell number in treated BCa cells. In a 2004 study, Chen W. *et al*. reported that concentrations of genistein ≥10μM inhibited the growth of MCF7 cells, an effect linked to alterations in the tyrosine kinase signaling pathway downstream of growth factor receptors [12]. In contrast, Li Z. *et al*. showed that even lower doses of genistein, as low as 5μM, were sufficient to reduce proliferation and induce apoptosis in MDA-MB-231 cells through the inhibition of the MAPK pathway [36]. Consistent with these findings, we observed that genistein concentrations similar to those reported by the referred authors also decreased cell viability in both MCF7 and MDA-MB-231 cells. Importantly, all genistein concentrations tested exhibited synergistic interactions with the chemotherapy agents evaluated in our study.

Clonogenic and scratch assays are commonly used to evaluate cancer cell proliferative and migratory capacities, respectively. While the clonogenic assay measures a cell’s ability to form colonies, reflecting its long-term survival and growth potential [37], the scratch assay assesses cell migration by creating a "wound" in a monolayer and observing how quickly cells close the gap [38]. In 2006, Raffoul J. *et al*. reported that in the prostate cancer cell line PC3, 15μM genistein, in combination with radiation, inhibited colony formation more effectively than either genistein or radiation alone [39]. Similarly, Fan P. *et al*. demonstrated that genistein at doses up to 15μM strongly reduced the clonogenic capacity of MCF7 cells [34]. Notably, the doses used in these studies align with the IC_10_ value for MCF7 cells observed in our work. However, we found that the MDA-MB-231 cell line was more susceptible to genistein’s anti-clonogenic effects than MCF7 cells. In 1999, Li Y. *et al*. showed that low doses of genistein induced apoptosis in MDA-MB-231 cells by downregulating BCL-2 and promoting the expression of the cell cycle inhibitor p21^WAF1^, while downregulation of p53 further reduced cell survival [40]. BCL-2 upregulation has been associated with enhanced clonogenic capacity in HeLa cells [41], and loss of p21^WAF1^ has been linked to increased clonogenic growth in keratinocytes [42]. Thus, the anti-clonogenic effects of genistein in MDA-MB-231 cells may be due to its ability to promote apoptosis and regulate cell cycle progression. However, further studies are needed to confirm this mechanism. In the scratch assay, genistein has been shown to significantly inhibit cell migration in various cancer cell lines. Specifically, 20-40μM genistein reduced the migration rate of HeLa cells [43], while 100 μM inhibited migration in A549-derived lung cancer stem-like cells [44]. However, studies in BCa cells have yielded conflicting results. A 2023 study found that 10μM genistein had no effect on the migration rate of MCF7 cells [45], while Ye D. *et al*. reported in 2017 that 20 and 40μM genistein suppressed migration in MDA-MB-231 cells [46]. These discrepancies may be attributed to differences in the doses and time points used in our study compared to those of the previous reports.

Cancer cell survival and migration are closely linked to their metabolic reprogramming. Unlike healthy cells, malignant cells rely on altered metabolic pathways that enable their growth and persistence. The Warburg effect, first described in 1924, demonstrated that cancer cells increase glucose uptake and glycolysis to elevate ATP production, even in the presence of sufficient oxygen for oxidative phosphorylation (OXPHOS). However, it is now widely accepted that cancer cells do not rely on glycolysis alone for energy production, with many also depending heavily on OXPHOS for ATP generation [47, 48].

In our evaluation of glycolytic activity and OXPHOS in breast cancer cells, we observed that genistein influenced the energetic phenotype differently in ER-positive and ER-negative cells. In MDA-MB-231 cells, genistein decreased the glycolytic reserve without affecting OXPHOS. In contrast, in MCF7 cells, although glycolysis remained unchanged, genistein treatment—either alone or in combination with chemotherapy—resulted in a decrease in ATP production. This is particularly noteworthy, as cancer cells typically exhibit upregulated mitochondria coupled to increased OXPHOS activity, which can dynamically change in response to anticancer therapy [48–50]. These results suggest that genistein alters cellular energy metabolism, leading to reduced ATP production capacity, particularly in MCF7 cells. Further investigation into mitochondrial fuel oxidation revealed that genistein alone did not significantly alter fuel utilization. However, in combination with chemotherapy, it reduced FA dependence and increased glucose dependence. In contrast, genistein had no effect on fuel dependence or capacity in MDA-MB-231 cells.

Genistein has been shown to modulate lipid metabolism, not only in healthy cells but also in cancerous cells. While its role in lipid metabolism across various cancers has been documented, there is currently no information on its effects in BCa. In the colon cancer DLD-1 cell line, genistein treatment (1 to 50 μM) reduced the expression of HMG-CoA reductase, an enzyme critical for *de novo* cholesterol synthesis and membrane formation in proliferating cells, in a dose-dependent manner [19]. In HT-29 human colon cancer cells, 200μM genistein inhibited lipid droplet aggregation by downregulating lipid metabolism markers such as perilipin-1, FASN, and PPAR-γ, an effect associated with apoptosis induction [51]. While studies directly linking genistein to lipid metabolism in cancer are limited, some research has explored its influence in combination with other lipids. For example, Horia and Watkins demonstrated in 2007 that combining 2.5μM genistein with the polyunsaturated FA docosahexaenoic acid reduced PGE_2_ and COX-2 expression in MDA-MB-231 cells, both of which are involved in inflammation and carcinogenesis [52]. Additionally, in LNCaP human prostate cancer cells, the combination of the cholesterol-lowering compound HPCD with escalating doses of genistein (up to 50μM) significantly decreased cell viability, promoted apoptosis, and inhibited AKT phosphorylation [53]. In HepG2 human hepatoma cells, 10μM genistein suppresses the expression of the master regulator of lipogenesis, SREBP-1, and its downstream effector FASN, while also inhibiting FASN enzymatic activity [54]. Similarly, genistein inhibited lipogenesis in H460 cells by reducing the expression and activity of SCD1 [31].

Our data demonstrate that genistein significantly reduced endogenous FAO in MCF7 cells, while in MDA-MB-231 cells, genistein alone increased FAO instead. Notably, in MCF7 cells, genistein treatment decreased CPT1 expression, which was associated with increased levels of saturated FAs, such as stearic and palmitic acid. Previous studies have shown that palmitate can impair mitochondrial function and induce apoptosis in BCa cells [55]. In contrast, in MDA-MB-231 cells, although no changes in CPT1 expression were observed, genistein treatment led to a reduction in stearic and palmitic acids. Additionally, pAMPK levels were decreased in both cell lines, regardless of the treatment. Phosphorylated AMPK typically promotes catabolic processes to generate ATP, while inhibiting anabolic pathways to conserve energy [56]. Thus, targeting AMPK activation is of interest, as its inactivation has been associated with anticancer effects [57]. In MCF7 cells, treatment with tamoxifen, or its combination with genistein, resulted in increased expression of SRBP1 and FAS, which may represent a compensatory response to the reduced FA utilization, potentially aimed at maintaining lipid biosynthesis. The reduced FAO, combined with an imbalance in lipid metabolite levels, may contribute to the establishment of a senescent phenotype, as observed in our genistein-treated cells [58]. However, the effects of genistein on senescence appear to be model-dependent. In previous studies, genistein has been shown to protect against senescence by activating AMPK in other systems, such as vascular smooth muscle cells [59].

To the best of our knowledge, this is the first study reporting the effects of genistein on the lipid profile of both ER-positive and ER-negative BCa cells. However, further research is needed to comprehensively explore the lipidomic effects of genistein across different BCa subtypes. Additionally, future studies should include *in vivo* models to assess the therapeutic potential of genistein and evaluate its applicability in clinical settings for patients with BCa.

## 5. Conclusion

Our study provides evidence for the synergistic anti-cancer effects of low-concentration genistein in combination with conventional chemotherapy in MCF7 and MDA-MB-231 human breast cancer cells. This combination significantly reduced clonogenic capacity and inhibited cell migration, two critical factors in cancer progression. Mechanistically, genistein appears to exert its anti-cancer effects by diminishing oxidative phosphorylation, reducing reliance on FA metabolism, and increasing intracellular FA concentrations, which may contribute to the induction of cellular senescence. These findings suggest that genistein, when used in combination with chemotherapy, holds promise as a potential adjuvant in breast cancer treatment. Future studies should explore the therapeutic potential of this combination in clinical settings, as well as its application with other conventional anticancer agents.

## Abbreviations

BCa: breast cancer
CRC: colorectal cancer
FAO: fatty acid oxidation
FFAs: free fatty acids
FAs: fatty acids
HMG-CoA reductase: 3-hydroxy-methylglutaryl-coenzyme A reductase
ER: estrogen receptor
GT: genistein+tamoxifen
GD: genistein+docetaxel
ICs: inhibitory concentrations
CI: combination index
OCR: oxygen consumption rate
ECAR: extracellular acidification rate
PVDF: polyvinylidene difluoride membrane
ANOVA: analysis of variance
CPT1: carnitine palmitoyltransferase 1
MMP-9: matrix metalloproteinase-9
OXPHOS: oxidative phosphorylation

## Ethics statement

Not required.

## Author contributions

STC, AVC, MJ: Experimentation and data analysis; YAAC, LGN, OG, ACB, RMB: Experimentation; NT and ART: Writing-review and providing of research facilities; ASC: Conceptualization, experimentation, data analysis, supervision, writing-original draft and review. All authors have read and approved the final version of the manuscript.

## Data availability statement

All data generated or analyzed in this work are included in the published article.

## Acknowledgments

Authors would like to thank Dr. Alejandro Zentella-Dehesa for his kind donation of the BCa cells used in this study.

